# Current range of *Agrilus planipennis* Fairmaire, an alien pest of ash trees, in European Russia and Ukraine

**DOI:** 10.1101/689240

**Authors:** Marina J. Orlova-Bienkowskaja, Alexander N. Drogvalenko, Ilya A. Zabaluev, Alexey S. Sazhnev, Elena Y. Peregudova, Sergey G. Mazurov, Evgenij V. Komarov, Vitalij V. Struchaev, Vladimir V. Martynov, Tatyana V. Nikulina, Andrzej O. Bieńkowski

## Abstract

**Context:** The first detection of *A. planipennis* in European Russia was in Moscow in 2003, when it began to spread.

**Aim:** To determine the range of *A. planipennis* as of 2020.

**Methods:** In 2017-2020, our Russian-Ukrainian research team examined >7000 *F. pennsylvanica* trees and >2500 *F. excelsior* trees in 84 localities of European Russia, Ukraine and Belarus.

**Results:** The current range exceeds the area of Spain and includes the Luhansk region of Ukraine and 16 regions of ER: Belgorod, Bryansk, Kaluga, Kursk, Lipetsk, Moscow, Orel, Ryazan, Smolensk, Tambov, Tula, Tver, Vladimir, Volgograd, Voronezh, and Yaroslavl. *Agrilus planipennis* was not detected in Belarus. The overwhelming majority of the infestations were found on *F. pennsylvanica*. All known cases of infestation of the native species (*F. excelsior*) are from artificial plantings.

**Conclusion:** *Agrilus planipennis* will appear in other European countries soon and damage *F. pennsylvanica.* Further surveys are necessary to determine whether *A. planipennis* infests *F. excelsior* in forests.

## 1. Introduction

The emerald ash borer, *Agrilus planipennis* Fairmaire (Coleoptera: Buprestidae), is a devastating alien pest of ash trees in European Russia and North America (Baranchikov et al. 2008; Herms and McCullough 2014; Haack et al. 2015). It is included in the list of 20 priority quarantine pests of the EU (EU 2019). The native range of this wood-boring beetle occupies a restricted territory in East Asia (Orlova-Bienkowskaja and Volkovitsh 2018). In 2003, *A. planipennis* was first recorded in European Russia, namely, in Moscow, and a severe outbreak and quick spread of the pest began (Baranchikov et al. 2008; Haack et al. 2015). By 2013, the pest was recorded in 9 regions of European Russia: from Yaroslavl in the north to Voronezh in the south (Straw et al. 2013; Orlova-Bienkowskaja 2014a). A probabilistic model of spread made in 2017 showed that in the next few years, the range of *A. planipennis* in Europe could expand significantly, and the pest could appear in neighboring countries (Orlova-Bienkowskaja and Bieńkowski 2018a).

*Fraxinus pennsylvanica* Marsh. was introduced from North America and is widely planted in European Russia as an ornamental and landscape tree. It is highly susceptible to *A. planipennis* both in North America and Russia (Herms and McCullough 2014; Baranchikov et al. 2014). The overwhelming majority of detected infestations of ash trees in European Russia correspond to this North American ash species (Baranchikov et al. 2008; Straw et al. 2013; Orlova-Bienkowskaja 2014a, etc.).

The only native ash species in European Russia is *F. excelsior* L. It is susceptible to the pest (Baranchikov et al. 2014). However, it is still unknown whether it is highly susceptible (as are the ash species native to North America) or less susceptible (as are the ash species native to Asia). On the one hand, the only cultivar of *F. excelsior* tested in a naturally infested experiment in America, cv. *Aureafolia*, had similar long-term mortality as that of *F. nigra* and *F. pennsylvanica* (Herms 2015). On the other hand, current experiments have shown that saplings of *F. excelsior* are much less susceptible than *F. nigra* Marsh. to *A. planipennis* (Showalter et al. 2019). *Fraxinus excelsior* is rare in the Moscow region and other regions of northern and central Russia. Therefore, information about the infestation of *F. excelsior* by *A. planipennis* is scarce and refers only to urban plantings and artificial shelterbelts, where *F. excelsior* is planted together with *F. pennsylvanica* (Straw et al. 2013; Baranchikov 2018). There is no information about the impact of *A. planipennis* on *F. excelsior* in natural forests. Therefore, there is a great deal of uncertainty surrounding the likely future impact of *A. planipennis* on ash in European forests (Straw et al. 2013).

The main aim of our study was to determine the range of *A. planipennis* in Europe as of 2020. An additional aim was to obtain information about the current condition of *F. excelsior* in natural forest stands in some regions occupied by *A. planipennis.*

## 2. Material and methods

### 2.1. Collation of previous records of*A. planipennis*

We have studied the *A. planipennis* range in European Russia since 2013 and have carefully examined all records that have appeared in journal articles, conference papers, official quarantine documents and other sources. These records, which comprise all of the published data on localities where *A. planipennis* has been detected, are compiled in Table 1.

**Table 1.** Previously published records of *Agrilus planipennis* in European Russia.

| Region | Locality | Year of first detection | Sources of information |
| --- | --- | --- | --- |
| Moscow reg. | Moscow<br>(*A. planipennis* occurs all over the city) | 2003 | Baranchikov et al. 2008; Izhevskii and Mozolevskaya 2010; Straw et al. 2013 |
| Moscow reg. | Manikhino | 2006 | Baranchikov et al. 2008 |
| Moscow reg. | Serpukhov | 2009 | Baranchikov et al. 2010 |
| Moscow reg. | Mozhaisk | 2009 | Baranchikov et al. 2010 |
| Moscow reg. | Mytiszhi | 2009 | Baranchikov et al. 2010 |
| Moscow reg. | Pushkino | 2009 | Baranchikov and Kurteev 2012 |
| Moscow reg. | Zelenograd | 2011 | Orlova-Bienkowskaja 2014b |
| Moscow reg. | Kolomna | 2012 | Orlova-Bienkowskaja 2014a |
| Smolensk reg. | Localities along the Moscow-Vyazma road | 2012 | Baranchikov and Kurteev 2012 |
| Moscow reg. | Sergiev Posad | 2012 | Baranchikov and Kurteev 2012 |
| Ryazan reg. | Vysokoe | 2012 | Baranchikov 2013 |
| Tula reg. | Shchekino | 2013 | Straw et al. 2013 |
| Tver reg. | Zubstov | 2013 | Straw et al. 2013 |
| Tver reg. | Novozavidovskiy | 2013 | Straw et al. 2013 |
| Tver reg. | Emmaus | 2013 | Straw et al. 2013 |
| Moscow reg. | Uzunovo | 2013 | Orlova-Bienkowskaja 2014a |
| Moscow reg. | Klin | 2013 | Orlova-Bienkowskaja 2014b |
| Moscow reg. | Staraya Kupavna | 2013 | Orlova-Bienkowskaja 2014a |
| Voronezh reg. | Voronezh | 2013 | Orlova-Bienkowskaja 2014a |
| Tula reg. | Tula | 2013 | Orlova-Bienkowskaja 2014a |
| Kaluga reg. | Kaluga | 2013 | Orlova-Bienkowskaja 2014a |
| Tver reg. | Konakovo | 2013 | Orlova-Bienkowskaja 2014b |
| Orel reg. | Orel | 2013 | Orlova-Bienkowskaja 2014a |
| Yaroslavl reg. | Yaroslavl | 2013 | Orlova-Bienkowskaja 2014a |
| Tambov reg. | Michurinsk | 2013 | Orlova-Bienkowskaja 2014a |
| Vladimir reg. | Petushki | 2013 | Baranchikov 2013 |
| Orel reg. | Livny | 2013 | Rosselkhoznadzor 2014 |
| Moscow reg. | Solnechnogorsk | 2014 | Orlova-Bienkowskaja and Belokobylskij 2014 |
| Moscow reg. | Planernaya | 2014 | Orlova-Bienkowskaja and Bieńkowski 2016 |
| Moscow reg. | Povarovka | 2014 | Orlova-Bienkowskaja and Belokobylskij 2014 |
| Smolensk reg. | Semirechje | 2014 | Zvyagintsev et al. 2015 |
| Smolensk reg. | Andreykovo | 2014 | Zvyagintsev et al. 2015 |
| Tver reg. | Khiminstituta settlement | 2015 | Peregudova 2016 |
| Voronezh reg. | Talovaya | 2018 | Baranchikov et al. 2018a |
| Lipetsk reg. | Elets | 2018 | Baranchikov et al. 2018a |
| Smolensk reg. | Smolensk | 2018 | Baranchikov and Seraya 2018 |
| Tver reg. | Doroshiha railway station | 2018 | Peregudova 2019 |
| Voronezh reg. | Olkhovatskoe<br>Lesnichestvo | 2018 | Rosselkhoz nadzor 2019 |
| Bryansk reg. | Majskij Park | 2019 | Rosselkhoz nadzor 2019 |
| Belgorod reg. | Roven'ki | 2019 | Baranchikov and Seraya 2019 |
| Belgorod reg. | Eriomovka | 2019 | Baranchikov and Seraya 2019 |
| Belgorod reg. | Nagorie | 2019 | Baranchikov and Seraya 2019 |
| Belgorod reg. | Aidar | 2019 | Baranchikov and Seraya 2019 |
| Voronezh reg. | Novobelaya | 2019 | Baranchikov and Seraya 2019 |

### 2.2. Localities of the survey

In 2017, 2018 and 2019, we examined more than 7000 green ash trees (*F. pennsylvanica*) and more than 2500 European ash trees (*F. excelsior*) in 84 localities of European Russia, Belarus and Ukraine (Table 2, Figs 1, 2).

**Table 2.**
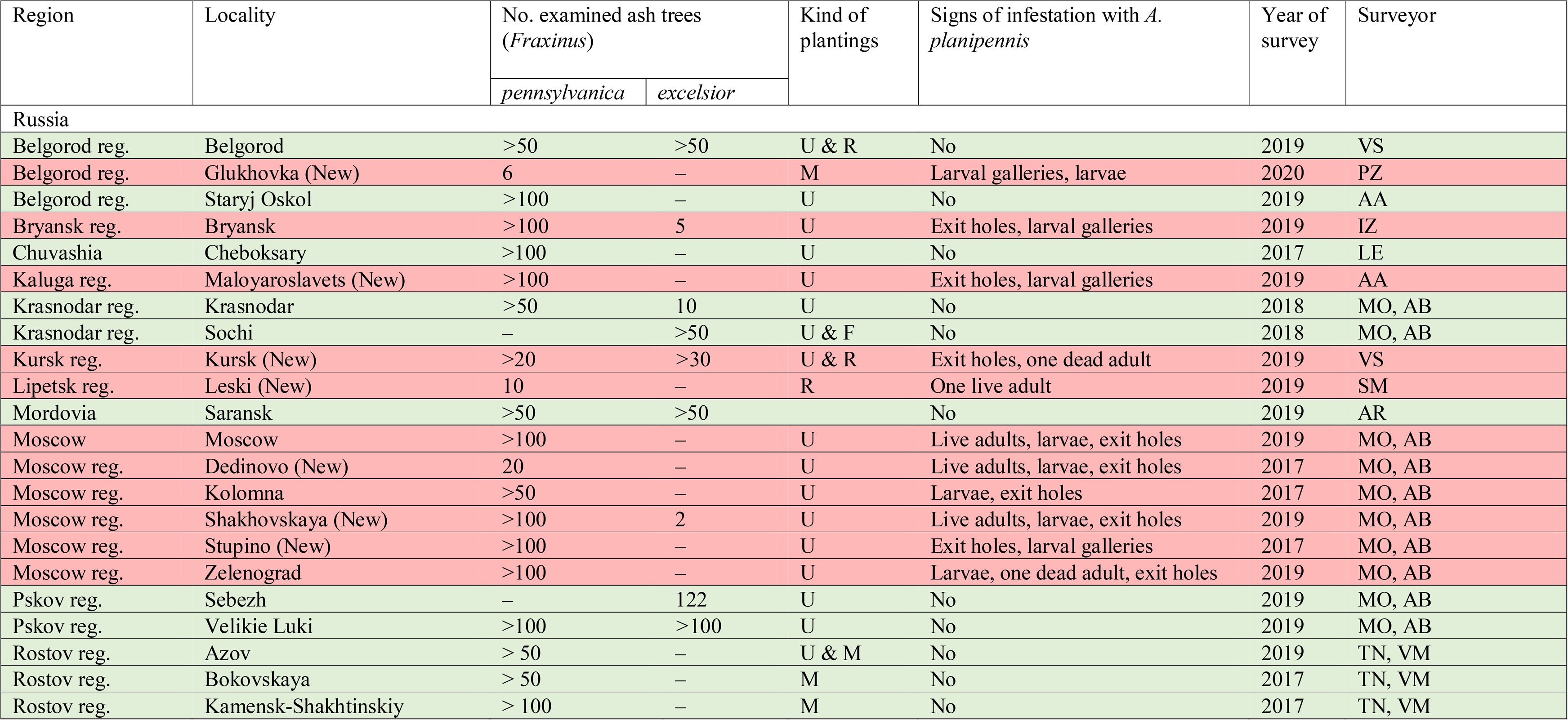
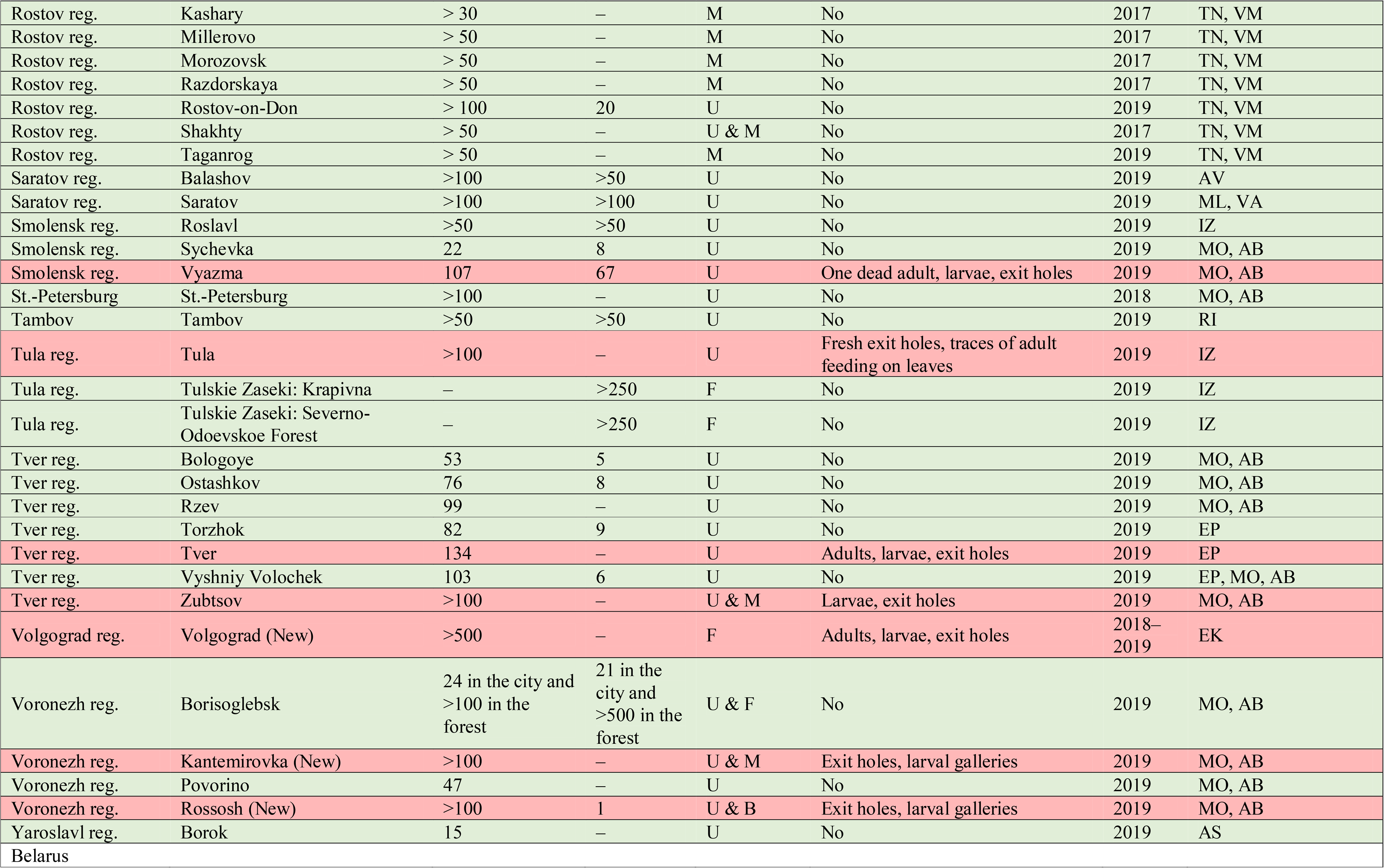
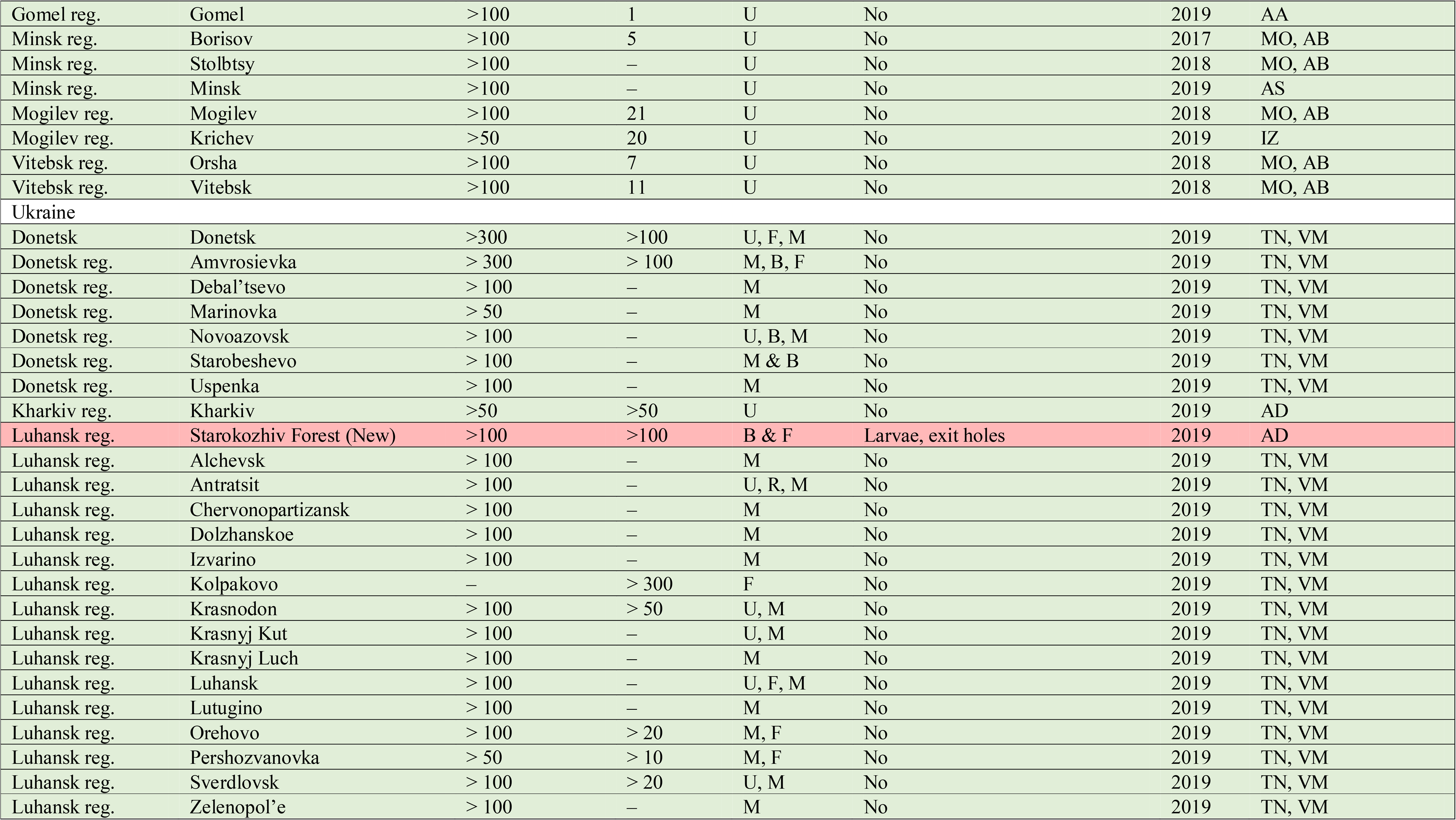
Surveys of ash trees in Russia, Ukraine and Belarus in 2017-2020. The localities where *Agrilus planipennis* was detected are marked with 490 red. The localities where *A. planipennis* was not detected are marked with green. New localities of detection are marked with the label “New”. Types of plantatings: F – forest, U – urban, M – along the motorway, R – along the railway, and B – shelterbelts. Surveys were made by AA – Andrzej A. Bieńkowski, AB – Andrzej O. Bieńkowski, AD – Alexander N. Drogvalenko, AR – Alexander B. Ruchin, AS – Alexey S. Sazhnev, AV – Alexey N. Volodchenko, EK – Evgenij V. Komarov, EP – Elena Yu. Peregudova, IZ – Ilya A. Zabaluev, LE – Leonid V. Egorov, ML – Mikhail V. Lavrentiev, MO – Marina J. Orlova-Bienkowskaja, PZ – Pavel A. Zavalishin, RI – Roman N. Ishin, SM – Sergey G. Mazurov, TN – Tatyana V. Nikulina, VA – Vasilij V. Anikin, VM – Vladimir V. Martynov, and VS – Vitalij V. Struchaev.

**Fig. 1.**
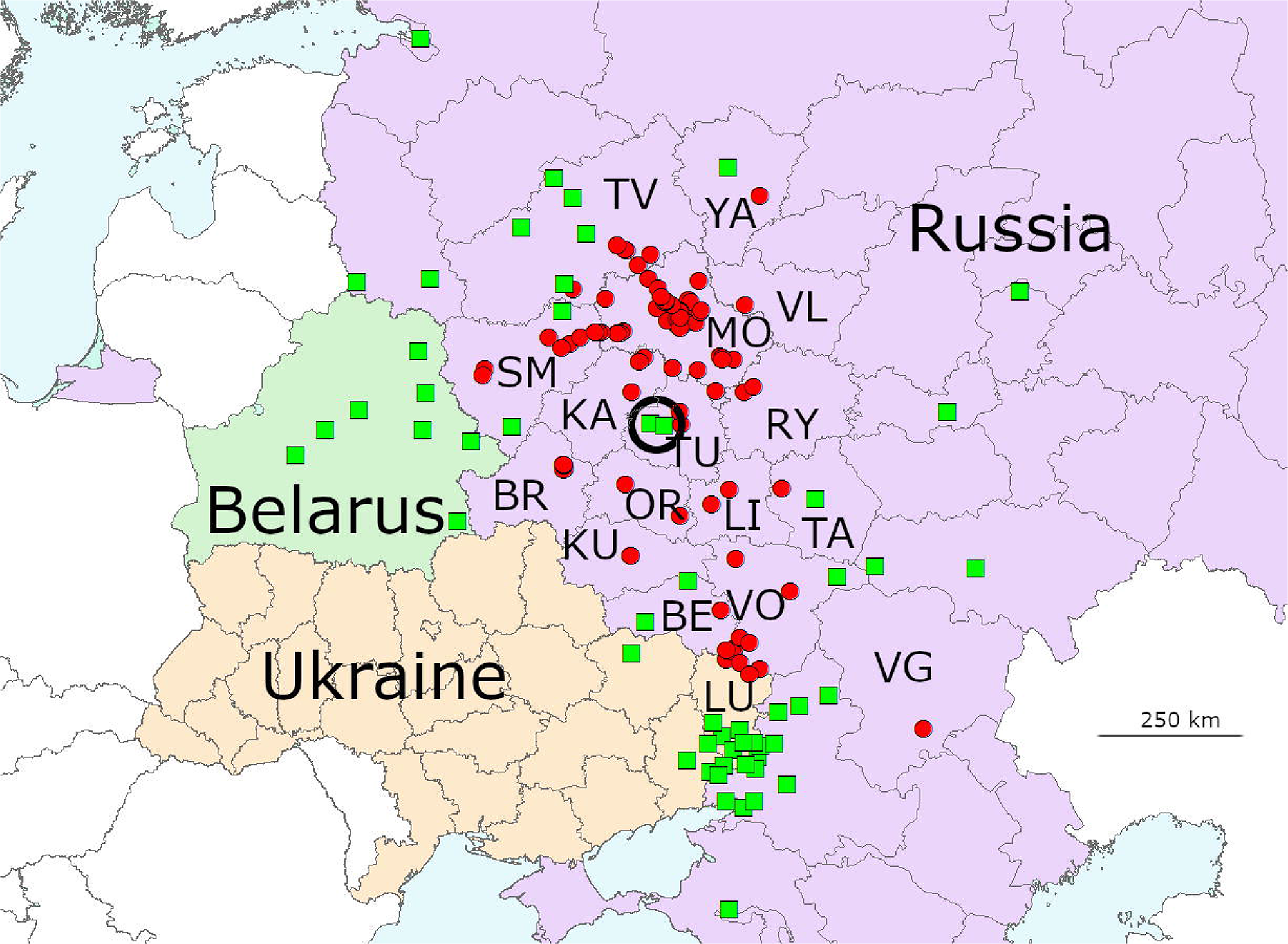
Range of *Agrilus planipennis* in European Russia (R) and Ukraine (U) in 2020. The red dots indicate the localities where *A. planipennis* was detected. The green squares indicate the localities where it was not detected during surveys in 2017-2019. The black circle indicates the localities of the surveys of *Fraxinus excelsior* in the broad-leaved Tulskie Zaseki Forest. BR – Bryansk region (R), BE – Belgorod (R), KA – Kaluga region (R), LI – Lipetsk region (R), LU – Luhansk region (U), MO – Moscow region (R), OR – Orel region (R), RY – Ryazan region (R), SM – Smolensk region (R), TA – Tambov region (R), TU – Tula region (R), TV – Tver region (R), VG – Volgograd region (R), VL – Vladimir region (R), VO – Voronezh region (R), and YA – Yaroslavl region (R). The sources are shown in Tables 1 and 2.

**Fig. 2.**
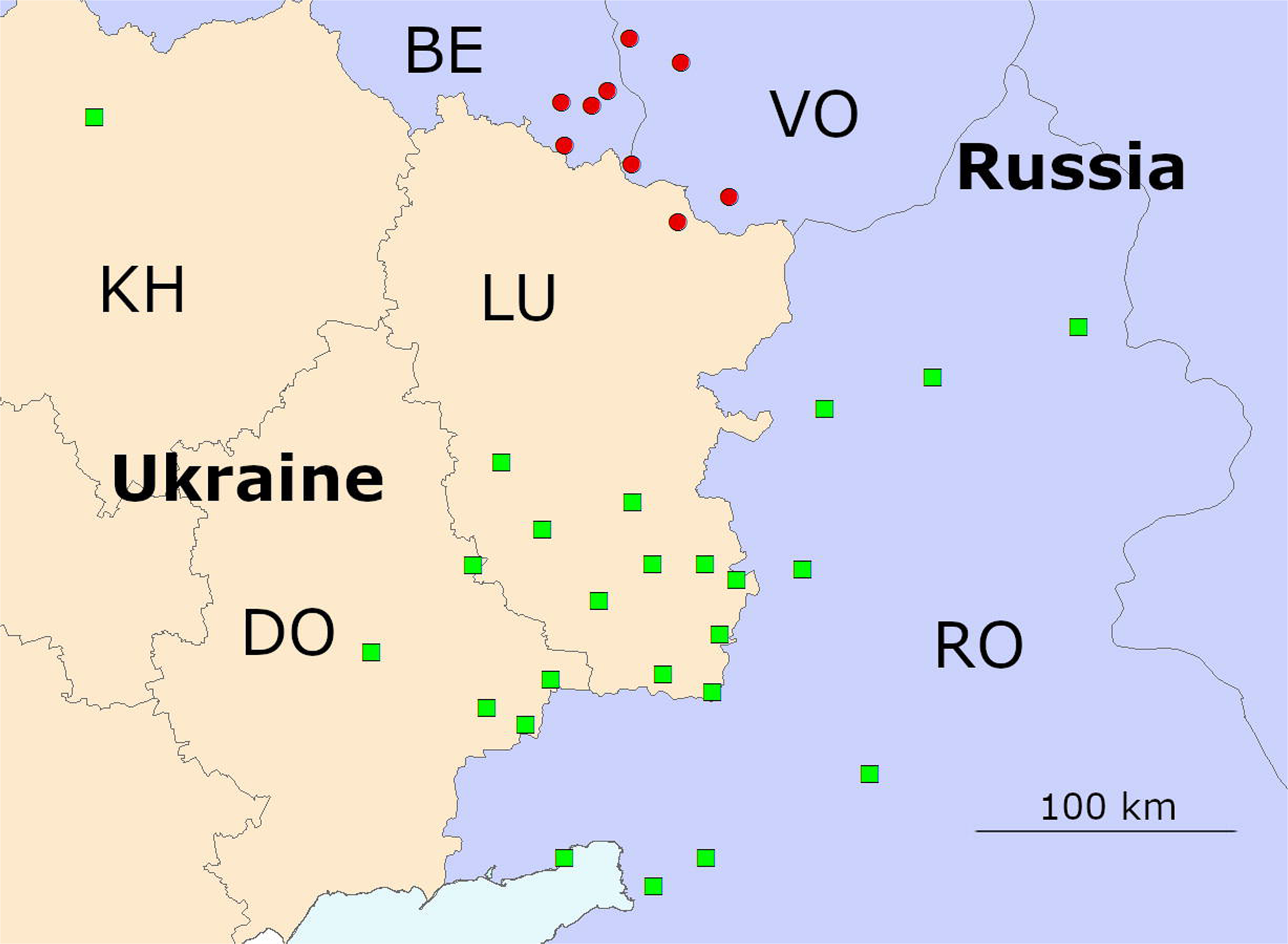
Results of the surveys in the eastern part of Ukraine (U) and the neighboring regions of Russia (R). The red dots indicate the localities where we detected *A. planipennis* in 2017-2019. The green squares indicate the localities where *A. planipennis* was not detected during surveys in 2017-2019. BE – Belgorod region (R), DO – Donetsk region (U), KH – Kharkiv region (U), LU – Luhansk region (U), RO – Rostov region (R), and VO – Voronezh region (R).

1. We examined ash trees in 7 Russian cities, where *A. planipennis* was found 4-16 years ago. The aim was to determine whether ash trees and populations of *A. planipennis* still existed there.
2. We examined ash trees in localities where *A. planipennis* had not been reported before. The aim was to determine the current range of the pest in Europe. We selected particular survey areas so that they were situated in all directions from the previously known range of *A. planipennis* in Europe. If we found the pest outside the previously known range, we conducted the next survey in a locality situated further in the same direction. We repeated this procedure until we found localities without the pest. This protocol was flexible since we used all possibilities to examine as many localities as possible.
3. We examined more than 500 *F. excelsior* in two localities in a large natural forest, Tulskie Zaseki, situated in the center of the current range of *A. planipennis* in European Russia. The aim was to determine whether there were trees infested with *A. planipennis* in this forest.

### 2.3. Method of survey

The survey of ash trees in each city always started from the main railway station. Usually, we stayed one day in each city and examined as many ash trees as we could. In most localities we examined artificial plantings in cities, along motorways and railroads and in shelterbelts. The survey of 500 *F. excelsior* trees in the large Tulskie Zaseki Forest was conducted in two localities situated 25 km apart (see Table 2).

We used the standard method of *A. planipennis* detection used by many authors (Baranchikov and Kurteev 2012; Straw et al. 2013, etc.). We looked for ash trees with symptoms of general decline (dieback of the upper part of the stem, reduced foliage, epicormic shoots, loose bark, etc.) (Fig. 3) and examined the lower part of the stem, below 2 m, for the presence of D-shaped exit holes of *A. planipennis* on the bark surface and larval galleries under the bark in the cambial region at the inner bark-sapwood interface (Figs 4, 5). If we found exit holes and larval galleries, we took photos and tried to collect larvae from under the bark and adults (live specimens on leaves or dead specimens in their exit holes). The larvae and adults were examined in the laboratory to confirm their identification and deposited in the authors’ collections. The following guides were used for identification: Chamorro et al. (2012) and Volkovitsh et al. (2019). The map was made with ArcView GIS 10.4.1 (Esri) in A.N. Severtsov Institute of Ecology an Evolution, Russian Academy of Sciences, Moscow (Esri Customer Number 282718).

**Fig. 3.**
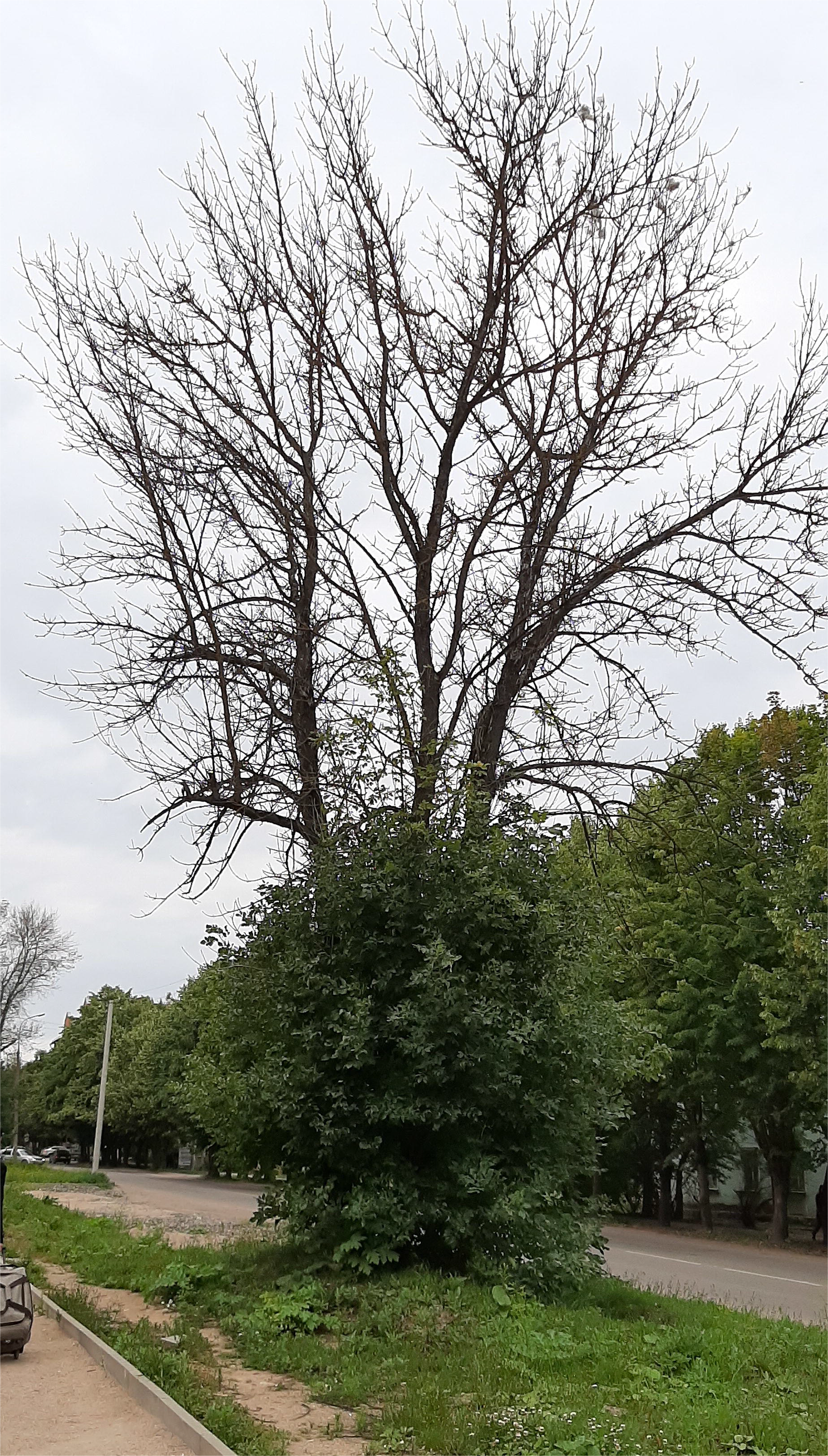
*Fraxinus pennsylvanica* tree heavily damaged by *Agrilus planipennis* in Vyazma, Russia in June 2019. Symptoms of general decline: dieback of the upper part of the stem and epicormic shoots. Photo by M.J. Orlova-Bienkowskaja.

**Fig. 4.**
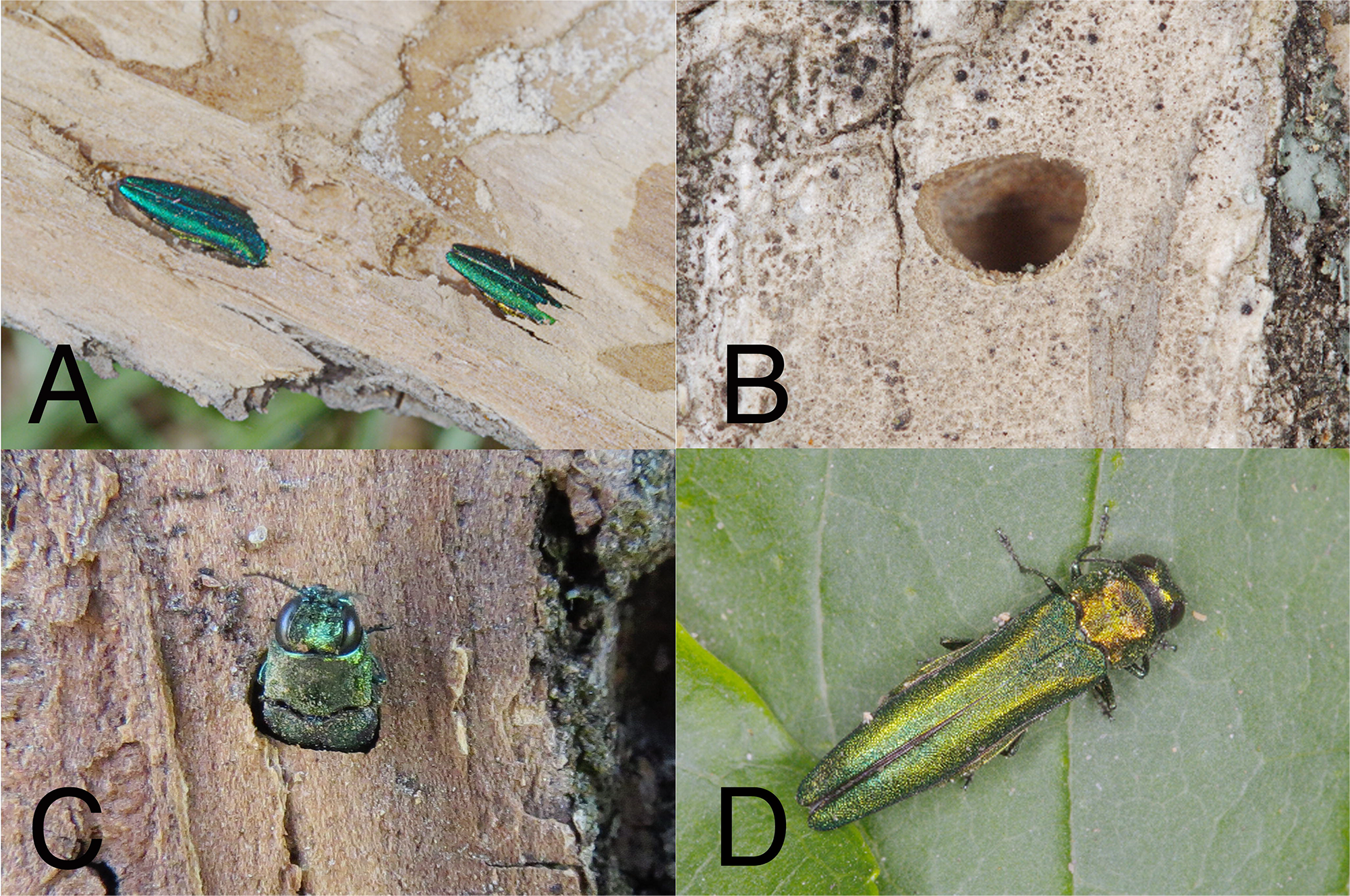
Adults of *A. planipennis* and their exit holes. **a.** Adults of *Agrilus planipennis* in pupal cells on *Fraxinus pennsylvanica* in Volgograd, Russia in 2018. Photo by E.V. Komarov. **b.** Exit hole of *Agrilus planipennis* on *Fraxinus pennsylvanica* in Volgograd, Russia in 2018. Photo by E.V. Komarov. **c.** Dead adult *Agrilus planipennis* found in Kursk in its exit hole on *Fraxinus pennsylvanica* in July 2019. Photo by V.V. Struchaev. **d.** Adult of *Agrilus planipennis* on a leaf of *Fraxinus pennsylvanica* in Volgograd, Russia in 2018. Photo by E.V. Komarov.

**Fig. 5.**
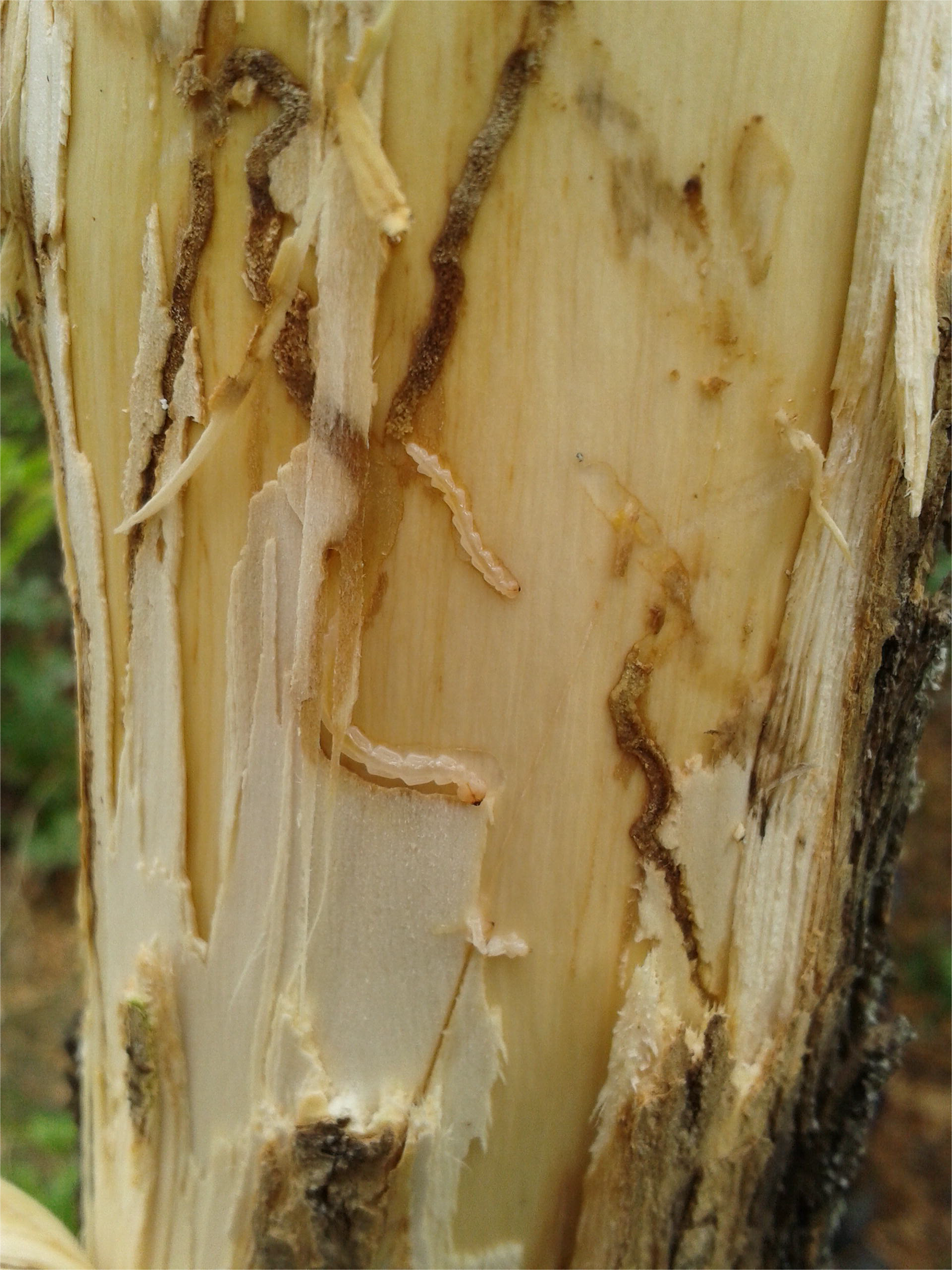
Larvae and larval galleries of *Agrilus planipennis* in *Fraxinus pennsylvanica* in the Luhansk region of Ukraine in September 2019. Photo by A.N. Drogvalenko.

## 3. Results

### 3.1. European Russia

#### 3.1.1. Localities with previous detections of *A. planipennis*

In 2019, we examined ash trees in 7 cities where *A. planipennis* was found 4-16 years ago: Moscow (first record in 2003), Kolomna (2012), Zelenograd (2011), Vyazma (2012), Tula (2013), Tver (2015) and Zubtsov (2013) (see Table 1 for information about the first records). The overwhelming majority of ash trees in these localities were *F. pennsylvanica*, which is one of the most common trees planted in the cities of European Russia. *Fraxinus excelsior* also occurred sometimes but much more rarely. In all these localities, ash trees were still common, and populations of *A. planipennis* still existed. Larvae or adults were found in all these localities in 2019.

Moscow was the entry point of invasion of *A. planipennis* in European Russia. By 2013, most ash trees in the city were heavily damaged (Orlova-Bienkowskaja 2014b), and some experts believed that ash trees would disappear throughout the city (Mozolevskaya 2012). However, surveys in 18 districts of Moscow in 2016 and 2017 showed that *F. pennsylvanica* was still common in the city, and *A. planipennis* had become rare (Orlova-Bienkowskaja and Bieńkowski 2018b). The survey in 2018 and 2019 confirmed this result. The reason for the decline in *A. planipennis* population in Moscow is unknown. It could be the result of mortality caused by the European parasitoid of *A. planipennis Spathius polonicus* Niezabitowski, 1910 (Hymenoptera: Braconidae: Doryctinae) (Orlova-Bienkowskaja and Belokobylskij 2014).

Infestations were found in all cities of the Moscow region surveyed by us in 2017-2019 (Table 2). In particular, a severe outbreak was recorded in 2019 in the city of Shakhovskaya. Almost all ash trees in the city were heavily infested. On 16 July 2019, more than 100 *A. planipennis* adults and larvae were collected there.

#### 3.1.2. Western regions: Tver, Pskov, Smolensk regions and St-Petersburg

We tried to find *A. planipennis* in 10 new localities of European Russia situated west and northwest of the border of the previously known range as of 2016: the Pskov region (Sebezh and Velikie Luki), Smolensk region (Roslavl and Sychevka), Tver region (Bologoye, Ostashkov, Rzev, Torzhok, and Vyshniy Volochek) and St. Petersburg. However, we detected no signs of the infestation with *A. planipennis* in these localities. It is interesting that we failed to find *A. planipennis* even in the city of Rhev, which is only 15 km northwest of Zubtsov, where an outbreak of *A. planipennis* was detected in 2013 (Straw et al. 2013) and is still continuing (based on our observations in 2019).

#### 3.1.3. Volgograd Region

In a planting of declining *F. pennsylvanica* trees (105 ha) on Sarpinsky Island in the Volga River (near the city of Volgograd) in October 2018 the trees were severely damaged by *A. planipennis*. The island is occupied by a floodplain forest, initially consisting of *Quercus*, *Populus*, and *Ulmus* trees, which was later replaced by an adventive plant: *F. pennsylvanica*. In different parts of the island, the infestation of ash trees varied from 20-30% to 90-100%. The decline in the ash plantings because of *A. planipennis* outbreak attracted the attention of the local television company (Vesti Volgograd 2018). In 2019, infested trees were also detected on the left bank of the Volga River, in the Volga-Akhtuba floodplain, on the Sarepta Peninsula and in the center of Volgograd city (Mira Street).

#### 3.1.4. Kursk region

In 2014, we surveyed more than 100 *F. pennsylvanica* trees in Kursk city near the main railway station and did not find evidence of *A. planipennis* (Orlova-Bienkowskaja and Bieńkowski, unpublished data). In 2016, the survey in Kursk was repeated by an expert from the All-Russian Center of Plant Quarantine, Y.N. Kovalenko, and he also did not find *A. planipennis* (unpublished data). In 2019, we examined more than 100 *F. pennsylvanica* trees in plantings along the railway near the main railway station in Kursk and found that at least half of the trees had *A. planipennis* exit holes. One dead adult specimen was collected in its exit hole. Additional examinations of *F. pennsylvanica* trees in the parks of Boeva Dacha and Pyatidesyatiletiya VLKSM showed that many trees were in poor condition and had *A. planipennis* exit holes.

#### 3.1.5. Lipetsk region

The Lipetsk region is surrounded by the Tula, Orel, Ryazan, Voronezh and Tambov regions, where *A. planipennis* was found in 2013 (Orlova-Bienkowskaja 2014a), but M.J. Orlova-Bienkowskaja did not find any trees infested with *A. planipennis* in the city of Gryazy in the Lipetsk region in 2013. However, many *F. pennsylvanica* trees were in poor condition, and the species *Agrilus convexicollis* Redtenbacher, which is oftenassociated with *A. planipennis*, was found (Orlova-Bienkowskaja and Volkovitsh 2014). In 2018, *A. planipennis* was first found in the Lipetsk region in the city of Elets (Baranchikov 2018). On 22 June 2019, we collected one adult *A. planipennis* approximately 40 km from Elets in plantings of *F. pennsylvanica* along the railway in Leski, Lipetsk region. Examination of 10 trees in this planting did not reveal exit holes, but all the trees of *A. planipenis* were in poor condition and had dieback in the upper canopy.

#### 3.1.6. Voronezh region

*Agrilus planipennis* was first detected in the Voronezh region in 2013 in the city of Voronezh (Orlova-Bienkowskaja 2014a). In 2018, it was detected in other localities: in the very south of the region (50°12'N, exact location is not known) and in the city of Talovaya (Baranchikov et al. 2018a), as well as in Olkhovatskoe Lesnichestvo (Rosselkhoznadzor 2019). In 2019, we surveyed ash trees in four other localities of the Voronezh region. *Agrilus planipennis* was not found in the northeast of the region (Borisoglebsk and Povorino), but was found in the following areas in the center and south of the region:

1. Rossosh. More than 100 heavily infested *F. pennsylvanica* trees with larval galleries and exit holes were found in a shelterbelt around the Chkalovskij district (50.201158 N, 39.604695 E). One *F. pennsylvanica* and one *F. excelsior* with exit holes and larval galleries were found near the main railway station (50.184266 N, 39.600577 E).
2. Kantemirovka. One *F. pennsylvanica* with *A. planipennis* exit holes and larval galleries was found in the territory of the House of Culture (49.698968 N, 39.860308 E). Trees of *F. pennsylvanica* with *A. planipennis* exit holes and larval galleries were also found near a gas station on the R-194 road (49.758194 N, 39.843341 E).

#### 3.1.7. Belgorod region

*Agrilus planipennis* was first detected in the Belgorod region in May 2019 in several localities close to the Voronezh region and the border of Ukraine (Baranchikov and Seraya 2019). Our surveys of ash trees in the cities of Belgorod and Staryi Oskol in summer 2019 did not reveal *A. planipennis*. In January 2020 larval galleries and larvae were found in the north-east of the Belgorod region (Table 2).

#### 3.1.8. Kaluga region

*Agrilus planipennis* was first detected in the Kaluga region in Obninsk city in 2012 and in Kaluga city in 2013 (Orlova-Bienkowskaja 2014a). A survey of more than 100 *F. pennsylvanica* trees in the city of Maloyaroslavets on 31 August 2019 revealed *A. planipennis* exit holes and larval galleries on at least 50 of the trees.

#### 3.1.9. Bryansk region

The first survey of ash trees in the city of Bryansk was conducted in 2013 and did not reveal signs of infestation by *A. planipennis* (Orlova-Bienkowskaja 2014a). In 2019, *A. planipennis* was detected in Majskij Park in Bryansk by the National Plant Protection Organization, and an official phytoquarantine zone was declared there (Rosselkhoznadzor 2019). On 1 July 2019, we examined ash trees in other districts of Bryansk and found that *F. pennsylvanica* trees in M.P. Kamozin Square and on Krasnoarmejskaya, Ulyaniva and Kharkovskaja streets were heavily infested with *A. planipennis*..

#### 3.1.10. Surveys of *F. excelsior* in the Tulskie Zaseki Forest in the center of the pest range in European Russia

The large broad-leaved Tulskie Zaseki Forest (65 000 ha) is situated in the Tula region in the center of the current range of *A. planipennis* in European Russia. *Fraxinus excelsior* is one of the main tree species of this forest. In 2013, *A. planipennis* was detected near this forest,in Tula and Shchekino as well as in the surrounding regions of Kaluga and Orel (Straw et al. 2013; Orlova-Bienkowskaja 2014a). *Fraxinus pennsylvanica* trees were very common in the cities of Tula and Shchekino, and almost all of them had been infested by *A. planipennis* (Baranchikov et al. 2018b and observations by I.A. Zabaluev). On 19-20 June 2019, more than 500 *F. excelsior* trees were examined in the Tulskie Zaseki Forest in two localities: near Krapivna village and in Severno-Odoevskoe Lesnichestvo. The trees were situated both along and within the forest. No signs of infestation by *A. planipennis* were found, and the trees appeared to be in good health.

### 3.2. Belarus

In 2017-2019, we tried to find *A. planipennis* in the eastern and central regions of Belarus in the cities of Borisov, Stolbtsy, Mogilev, Orsha, Vitebsk, Minsk, Krichev and Gomel. We did not find any signs of infestations despite *F. pennsylvanica* and *F. excelsior* being commonly planted in these cities (Table 2).

### 3.3. Ukraine

On 20-22 June 2019, ash trees in Starokozhiv Forest and field shelterbelts in its vicinity were examined (in the Markivka district of the Luhansk region of Ukraine). The forest consisted of *Acer platanoides* L., *F. excelsior*, *Pyrus* sp. and *Quercus* sp. The undergrowth and edges consisted of *Acer campestre* L., *Acer tataricum* L., *Fraxinus excelsior, F. pennsylvanica, Prunus fruticosa* Pallas. and *P. spinosa* L. The shelterbelts consisted mainly of *F. pennsylvanica.* During the examination of 250 ash trees, three *F. pennsylvanica* trees damaged by *A. planipennis* were detected. These trees were situated at the edge of the shelterbelts and had diameters of 7-10 cm. Characteristic D-shaped exit holes were situated at a height of 50-200 cm above the groundline. The infested trees had dieback of the upper branches, and foliage density was reduced (i.e., the trees had fewer and smaller leaves).

On 2 July 2019, we posted an early version of this manuscript on the Internet as a preprint (Orlova-Bienkowskaja et al. 2019). Immediately following the appearance of this preprint on the Internet, the National Plant Protection Organization of Ukraine conducted an official survey in the same area and did not detect *A. planipennis*. Since there were no specimens or photos for confirmation, our record of *A. planipennis* in Ukraine was considered unreliable (EPPO 2019a).

On September 4-6 2019, A.N. Drogvalenko visited the Markivka district in the Luhansk region of Ukraine again and repeated his survey of ash trees. He found the same three trees and more than 40 other *F. pennsylvanica* trees that were heavily infested with *A. planipennis*, took photos of larvae, larval galleries (Fig. 4) and exit holes and collected more than 20 larvae of different instars, including the last (4^th^) instar. The coordinates of these trees, from which the larvae were collected, are as follows: 49.614991 N, 39.559743 E; 49.614160 N, 39.572402 E; and 49.597043 N, 39.561811 E (roadside planting). The larvae were deposited in a collection in Kharkiv, Ukraine by A.N. Drogvalenko (Drogvalenko et al. 2019). After that, the presence of *A. planipennis* was also confirmed by the Ukrainian NPPO (EPPO 2019b). On October 22 2019, *A. planipennis* was found within 2 km radius from initial observation point outside the established quarantine zone (Meshkova 2019).

Our survey of ash trees in other localities of Ukraine (Luhansk region, Donetsk region and Kharkiv city) did not reveal any *A. planipennis* infestations despite both *F. pennsylvanica* and *F. excelsior* being usual in these regions and occurring in both natural forests and in plantings (shelterbelts and along roads) (Table 2, Fig. 2).

## 4. Discussion

### 4.1. Current borders of *A. planipennis*range in Europe

Previously published records (Table 1) and our surveys in 2017-2020 (Table 2) indicate that by 2020, *A. planipennis* occurred in the Luhansk region of Ukraine and in at least 16 regions of European Russia: Belgorod, Bryansk, Kaluga, Kursk, Lipetsk, Moscow, Orel, Ryazan, Smolensk, Tambov, Tula, Tver, Vladimir, Volgograd, Voronezh, and Yaroslavl (Fig. 1). It should be noted that these results were obtained by simple surveys of ash trees without the use of pheromone traps. Usually, the signs of *A. planipennis* become visible only several years after the establishment of this pest in a region (Haack et al. 2015). Therefore, the real range of *A. planipennis* in Europe could be even more extensive.

The westernmost known *A. planipennis* localities are in Semirechje (Zvyagintsev et al. 2015) and Smolensk city (Baranchikov and Seraya 2018) in the Smolensk region. In the northwest, the border of the range is near Zubtsov and Tver in the Tver region. The northernmost locality is Yaroslavl. The eastern border of the range in European Russia is poorly known. The extreme localities in this direction are Yaroslavl, Petushki (Vladimir region), Vysokoe (Ryazan region), Michurinsk (Tambov region) and Volgograd. However, it is unknown if this is the true eastern border of the range, since few surveys were conducted further east of these localities. The most eastern and the most southern known locality was Volgograd. In the southwest, the known border crossed the Bryansk, Kursk, Belgorod and Voronezh regions of Russia and the Luhansk region of Ukraine. In the southwest, the border of *A. planipennis* known range almost coincided with the Russian-Ukrainian border. Therefore, it is quite possible that *A. planipennis* will soon be found on the other side of the state border, in the Chernihiv, Sumy and Kharkiv regions of Ukraine.

### 4.2. Dynamics of the range since 2013

The northern and northwestern borders of *A. planipennis* range in European Russia in 2019 were almost the same as those in 2013. The northernmost locality–Yaroslavl–is at a latitude of 57.63 N, i.e., much further north than the northernmost locality of *A. planipennis* in North America (47.31 N) (Emerald Ash Borer Info 2019) and in Asia (49.42 N) (Orlova-Bienkowskaja and Volkovitsh 2018). Therefore, it is unknown whether *A. planipennis* can spread further north to St. Petersburg and northern Europe or if it has already reached its potential border in the north. To answer this question, an ecological model of the potential range of *A. planipennis* in Europe should be created, taking into account that the generation time in Moscow is 2 years (Orlova-Bienkowskaja and Bieńkowski 2016).

In the south, the situation is much worse. *Agrilus planipennis* is quickly spreading. It has now appeared in Ukraine and in the nearby areas of European Russia, such as the Volgograd, Bryansk, Kursk, Belgorod regions and the south of the Voronezh region.

### 4.3. Perspectives for European forestry

The distance from Moscow to Volgograd is more than 900 km, which is more than the distance from Moscow to Lithuania, Latvia and Estonia. Obviously, the pest can be detected in any part of Eastern Europe. In addition, *A. planipennis* could soon appear in Kazakhstan, given that the distance between Volgograd and the border of Kazakhstan is approximately 150 km.

Since there are no infested *F. excelsior* trees in the large Tulskie Zaseki Forest in European Russia, it seems that *F. excelsior* could be more resistant to the pest than ash species native to North America are, at least in natural forest stands. All cases of infestation of *F. excelsior* in Russia correspond to trees that grew near plantings of *F. pennsylvanica*. *Fraxinus pennsylvanica* is highly susceptible to *A. planipennis* and trees of this species are a source for the outbreak of the pest, which can also attack *F. excelsior* trees growing nearby. A similar situation was observed in China; in 1964, plantings of *Fraxinus americana* L. introduced from North America triggered the outbreak of *A. planipennis* in Harbin (Liu et al. 2003). The pest damaged both *F. americana* and native ash species in the region, but after the *F. americana* trees were cut, the outbreak ended.

*Agrilus planipennis* has not yet become a major forest pest in European Russia or Ukraine. It occurs in artificial plantings, such as urban plantings, along roads, railroads and in shelterbelts. The overwhelming majority of infested and dead trees are *F. pennsylvanica*. The only ash species native to European Russia, *F. excelsior*, is also susceptible to *A. planipennis*, but there is still no evidence of the widespread mortality of this ash species in European forests. There is no doubt that *A. planipennis* will severely damage plantings of *F. pennsylvanica* in Europe. However, the prognosis for the future of *F. excelsior* in European forests is still uncertain.

## 5. Conclusion

In just 16 years, following the first record in Europe, *A. planipennis* has spread over an area of 600 000 km^2^ (i.e., its current range exceeds the area of Spain). The range includes 16 regions of European Russia and the Luhansk region of Ukraine. The pest is quickly spreading to the south and will undoubtedly appear in other European countries soon. No expansion of the range to the north has been detected since 2013. No case of serious *A. planipennis* damage to *F. excelsior* in European forests has been detected yet. Therefore, it is still unknown whether *A. planipennis* will become a devastating forest pest in Europe or just a pest of urban plantings. An extensive survey of the European ash (*F. excelsior*) in natural forests is necessary to determine the future possible impact of *A. planipennis* on European forests. We hope that this article will encourage experts from different European countries to study *A. planipennis* before this pest reaches the EU.

## Acknowledgments

We thank A.B. Ruchin from Mordovski Nature Reserve, L.V. Egorov from Prisurski Nature Reserve, A.N. Volodchenko from Saratov State University, A.A. Bieńkowski from Moscow State University, R.N. Ishin from Tambov State University, P.A. Zavalishin from Saint Petersburg State Forest Technical University and V.V. Anikin and M.V. Lavrentiev from Saratov State University for the information about the surveys of ash trees in different regions of European Russia and Belarus and S.A. Bieńkowski for the preparation of the maps and photos.

## Notes

#### Summary of Updates

1. Immediately following the appearance of the first version of this preprint on the Internet, the National Plant Protection Organization of Ukraine conducted an official survey in the same area and did not detect A. planipennis. Since there were no specimens or photos for confirmation, our record of A. planipennis in Ukraine was considered unreliable (EPPO 2019a). On September 4-6 2019, A.N. Drogvalenko visited the Markivka district in the Luhansk region of Ukraine again and repeated his survey of ash trees. He found the same three trees and more than 40 other F. pennsylvanica trees that were heavily infested with A. planipennis, took photos of larvae, larval galleries and exit holes and collected more than 20 larvae of different instars, including the last (4th) instar. After that, the presence of A. planipennis was also confirmed by the Ukrainian NPPO (EPPO 2019b). 2. Six month have passed after the first submission of this article to the journal. And a lot of new localities of A. planipennis were found during this time. We have added all these new data to the current version manuscript. The most important findings are in Kursk and Belgorod Regions of Russia.

